# Transcriptome-wide analysis of primary human endothelial cell responses to 1 hour of protein translation inhibition identify nonsense mediated decay targets and a non-coding *SLC11A2* exon as an acute biomarker

**DOI:** 10.1101/2023.09.21.558767

**Authors:** Adrianna M. Bielowka, Fatima S. Govani, Dilip Patel, Maria E Bernabeu-Herrero, Dongyang Li, Micheala A. Aldred, Inês G. Mollet, Claire L Shovlin

**Author notes:** Correspondence:* Claire L. Shovlin PhD FRCP, National Heart and Lung Institute, Imperial Centre for Translational and Experimental Medicine, Imperial College London, Hammersmith Campus, Du Cane Road, London W12 0NN, UK.

## Abstract

Nonsense mediated decay (NMD) lowers the cellular concentration of spliced RNAs harboring premature termination codons (PTC), and inhibition has been proposed as a potential therapeutic method. Conversely, NMD plays regulatory roles throughout the eukaryotic kingdom, including when protein translation is inhibited acutely as part of the integrated stress response. To define tools for endothelial evaluations of therapeutic NMD inhibition, and quantification of subtle cellular stress states, natural endothelial-expressed targets were examined via whole transcriptome RNA sequencing of primary human microvascular endothelial cells (HMECs) treated for 1h with cycloheximide, a protein translation and NMD inhibitor. Genes differentially expressed after 1h cycloheximide overlapped with genes differentially expressed many days after NMD-specific knockdown in other cell types. For endothelial cells, customized novel scripts used 255,500 exons in media-treated HMEC and 261,725 exons in cycloheximide-treated HMEC to predict 1h cycloheximide-stabilized exons. RT-PCR and RNASeq validations in other endothelial cells highlighted exon 3B of the iron transporter *SLC11A2* (also known as *NRAMP2/DMT1*) as a novel exon in a transcript most consistently stabilized. Exact junctional alignments to *SLC11A2* exon 3B were confirmed in blood outgrowth endothelial cells (BOECs) from 3 donors at mean 5.9% (standard deviation 2.0%) of adjacent constitutive exon expression, increasing 3.7-fold following 1h treatment with cycloheximide. Relevance beyond endothelial cells is supported by *SLC11A2’s* wide expression profiles, genome-wide associations with microcytic anemia, biomarker status for poor prognosis ovarian cancer, and exon 3B sequence in RefSeq non-coding transcript NR_183176.1. The studies contribute understanding to functions affected acutely by NMD/translation inhibition and provide a stimulus for further studies in experimental, stress, and therapeutic settings.

## INTRODUCTION

Endothelial cells, as for all other eukaryotic cells, employ nonsense mediated decay (NMD) as one of the major physiological processes to promote translational fidelity, and provide RNA-based responses to triggers that would otherwise perturb cellular homeostasis. [1,2] NMD plays regulatory roles throughout the eukaryotic kingdom, is translation dependent, and reversed by broad protein synthesis inhibitors such as cycloheximide,[3] in addition to more NMD specific regulators.[4–7] NMD is blunted by a number of stressors including endoplasmic reticulum stress when unfolded or misfolded proteins are accumulating, and exogenous stress when NMD is downregulated as a consequence of stress signals that inhibit protein translation.[8–10] In turn, the integrated stress response which is triggered by multiple stresses such as reactive oxygen species (ROS) generation, hypoxia, and limited amino acid supply, depends upon key natural NMD targets (ATF3, ATF4 and CHOP) that provide mechanisms to augment cellular stress responses.[9–11]

NMD predominantly takes place while a newly synthesised and spliced mRNA remains bound by the CBP20-CBP80 cap-binding protein (CBP) heterodimer, during the first (“pioneer”) round of protein translation. During pre-mRNA splicing, exon junction complexes (EJCs) of proteins are deposited upstream of each exon-exon junction and are displaced as the ribosome moves along the transcript.[1,2,12] Since natural stop codons are usually located in the terminal exon, once protein translation is completed, all EJCs will have been removed. However, if a premature stop codon (PTC) is present upstream of the last exon-exon boundary, translation will terminate early, leaving at least one EJC bound to the mRNA, and unless within 50-55 nucleotides of the PTC, the EJC triggers NMD.[1,2,12] EJC-independent NMD also occurs, and can target cytoplasmic mRNAs in which the CBP20/CBP80 heterodimer has been displaced by eukaryotic initiation factor 4E (eIF4E).[1,2] NMD depends upon the highly conserved ATP-dependent RNA helicase UPF1,[4,7] with activity moderated by varying NMD factors in different settings.[1,2,4–8] NMD is illustrated in a simplified manner in Figure 1A.

**Figure 1:**
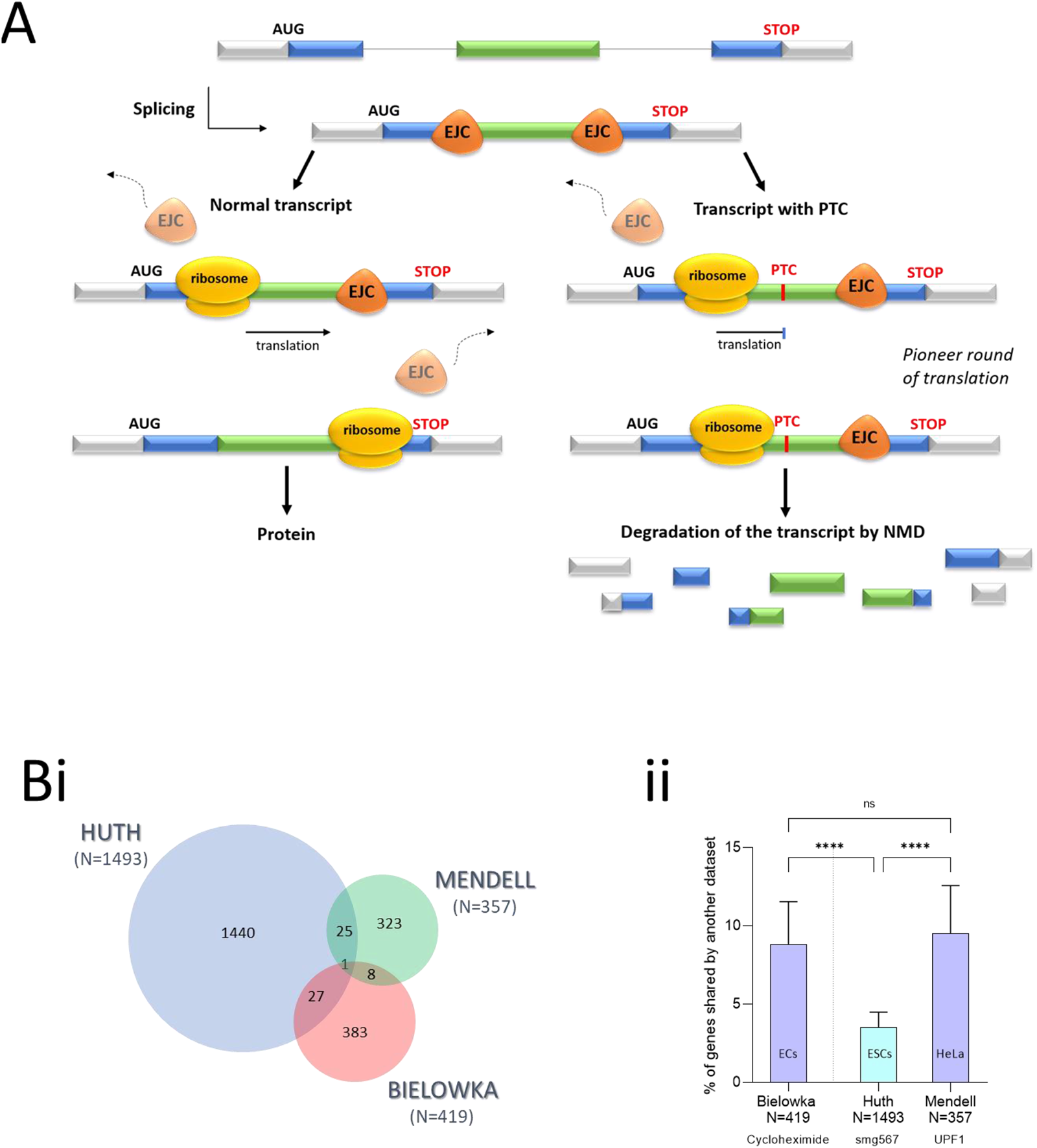
NMD and validations. **A) Cartoon of premature termination codon (PTC) recognition and activation of nonsense mediated decay (NMD).** Exon junction complexes (EJCs) comprising multiple splicing factors, mRNA export proteins and core EJC components[12] are deposited at exon-exon junctions during pre-mRNA splicing, and displaced as the ribosome moves along the transcript. Left: normal transcript with a terminal exon natural stop codon (UGA, UAA or UAG): On completion of protein translation, all EJCs will have been removed. Right: transcript with a PTC: Translation will terminate early, leaving EJCs bound to the mRNA, and if sited at least 50-55 nucleotides upstream of the final exon-exon junction, this triggers NMD. For details, the reader is referred to recent biochemical reviews.[1,2] **B) Comparisons of the proportions of genes identified as differentially expressed in 3 studies of NMD inhibition. *i*)** Scale representations of number of genes and overlaps in current study, by Huth et al [34] and by Mendell et al.[35] Note the different timescales of 1hr (current study), 72hr [35] and >12 days.[34] ***ii*)** Percentages of genes identified as differentially expressed in each of the 3 studies that were also identified as differentially expressed in one other study. Kruskal Wallis p-value <0.0001, Dunn’s post test,**** p<0.0001. EC endothelial cells (human); ESC, embryonic stem cells (mouse); HeLa (human).

Alongside fidelity control mechanisms, and stress responses, NMD is a major contributor to post transcriptional gene regulation. An estimated one-third of alternatively spliced transcripts harbour a PTC, and the term regulated unproductive splicing and translation (RUST) has been used to describe regulation of gene expression through the coupling of NMD and alternative splicing.[13] Up to 50% of human transcripts contain at least one upstream open reading frame (uORF) which contains a PTC that can trigger NMD when the uORF is used.[14] New data highlight a role for NMD in circadian clock regulation of gene expression,[6] and further examples are described in recent data manuscripts and reviews.[1,2,4–7] Thus, the cellular environment differs according to the degree to which NMD is operational, and biomarkers of such a state would be valuable for experimental cell biologists.

NMD is considered a therapeutic target where aberrant premature termination codons (PTCs) are generated due to somatic or germline pathogenic DNA sequence variants that cause disease.[15–17] In conjunction with nonsense readthrough approaches, NMD inhibition is attractive since available transcript concentration would otherwise be a major limitation.[17] NMD inhibition has also been proposed for states where tumors use NMD to downregulate tumor suppressor genes.[18,19] Nevertheless, since inhibition of nonsense mediated decay is a nonspecific strategy, it can potentially affect any transcripts harboring a PTC. Further, while PTC+ mRNAs are default targets of NMD, recent findings indicate that mRNAs with features other than PTCs are also major contributor to biology consequences when NMD is inhibited. NMD inhibition would therefore stabilize not only mRNAs with nonsense mutations, but also natural targets of NMD, some of which could have detrimental effects on the cell.

Here we report experimental data that provide further insight into the effect of protein translation and NMD inhibition on endothelial physiology and assays for future studies, using a brief 1h treatment with cycloheximide to mimic an acute cellular response to stress.

## MATERIALS AND METHODS

### Endothelial cell viability in the presence of cycloheximide

Since prolonged cycloheximide treatment results in cell death, treatment times in specific primary endothelial cell types were optimized in pilot experiments, during experimental design. Using concentrations already optimized for inhibition of NMD,[3,20–24] primary human microvascular endothelial cells (HMEC) and primary human pulmonary artery EC (HPAEC, Promocell (PromoCell GmbH, Heidelberg)) were cultured in antibiotic-free Promocell media (PromoCell GmbH, Heidelberg) for which supplements included 5% fetal calf serum (FCS) as recommended for microvascular EC, and 2.5% FCS for large vessel EC.[24] For experimental treatments of confluent EC, the appropriate media was replaced with fresh, prewarmed EC-specific media with and without cycloheximide 100μg/mL (Sigma-Aldrich, US), and EC cultured for a further 1h or 3h before imaging.

### Primary human microvascular endothelial cell RNASeq

Initial RNASeq evaluations were performed in normal human dermal microvascular EC (HMEC) purchased from Promocell (PromoCell GmbH, Heidelberg). HMEC lot number 0020208.1 was isolated from facial skin of a 63 year old female Caucasian with 89% viability, a population doubling time of 26.6hs, and free of bacterial, fungal, mycoplasma, HIV-1 or HBV/HCV infection (Certificate of Analysis). HMEC culture was as described previously,[15,24,25] in antibiotic-free Promocell media. For the experimental treatments, replicate wells were treated with control media or media containing cycloheximide 100μg/mL for 1h to inhibit protein translation and abrogate NMD.[3,20–24]

Ribosomal (r)-RNA-depleted total RNA was used to prepare strand-specific whole transcriptome libraries and clusters generated for short single-read standard sequencing on an Illumina platform as described elsewhere.[24] 26 million 40nt strand specific valid reads were obtained from each sample, defined by Phred scores ≥20 and a maximum of 2 mis-matches per read aligned to NCBI36-genome assembly hg18.[24]

The number of sequenced reads aligning to microRNA stem-loop sequences from miRBase were counted using custom Perl scripts, and normalized to the total number of valid reads and exon size.[24] Valid reads were aligned to spliced transcripts from ExonMine[26] using Seqmap.[27] Differences in gene expression between two samples were evaluated using read counts for all exons and junctions for the gene strand only. From 15,756 alignments, the 419 genes with the most differential alignments after 1h treatment with cycloheximide or fresh media were clustered using the Database for Annotation, Visualization and Integrated Discovery (DAVID [28,29]). To control for bias, clustering was also carried out on a random list of 419 genes generated from the whole cycloheximide RNAseq dataset. For each gene cluster, the most significant processes with the smallest Bonferroni p-value were selected.

Separately, scripts were written to select pre-miRNAs that demonstrated alignments at least 20% higher in cycloheximide-treated cells or in media-treated cells. Target genes of identified pre miRNAs were analysed using TargetScan.[30,31] Genes were sorted depending on the number of miRNA binding sites and a maximum of 3000 targets were used for each of the miRNAs for Gene Ontology Clustering to identify associated cellular processes.

### Validations compared to other methods of NMD inhibition

Cycloheximide inhibits NMD by inhibiting eukaryotic translation elongation, binding to the 60S ribosomal subunit E-site.[32,33] To examine relevance of findings specific to NMD inhibition, genes identified as differentially expressed in endothelial cells treated with cycloheximide for 1h were compared to data published in earlier studies using NMD-specific inhibition methods. We focused on the data from knockdown of NMD factors SMG5, SMG6 or SMG7 in mouse embryonic stem cells [34] and 72h siRNA inhibition of UPF1 in HeLa cells.[35]

### Validations in other cell types

Initial validations were performed in human umbilical vein endothelial cells (HUVECs) purchased from Promocell (PromoCell GmbH, Heidelberg), and cultured in antibiotic-free EGM-2 medium (Lonza, US) supplemented with 10% fetal calf serum (FBS) (Lonza, US). Quantitative reverse transcription PCR (RT-qPCR) was performed as described.[24] Briefly, forward and reverse oligonucleotide primers were designed to target gene exon-exon junctions to limit genomic DNA amplification. Amplification conditions were first optimized, performed in 20µl reaction volumes using 3 min at 95⁰C, and then 40 cycles of 10 sec at 95⁰C and 30 sec at 60⁰C, before size separation of 1/10 volume by agarose gel electrophoresis. Where multiple bands indicated off-target binding, approaches to increase specificity of the oligonucleotide primers were required, particularly increasing the annealing temperature until a single band was observed.

Further validations were performed in blood outgrowth endothelial cells (BOECs) established from healthy volunteers using methods as described,[15,36] with ethical approval from the East of Scotland Research Ethics Service (16/ES/0095). All participants gave written informed consent. All EC were from separate donors, not used beyond passage 5, and allowed to reach confluence before treatments. HUVEC and BOEC cells were plated at 4 x 10^5^ in a 6 well place and incubated overnight at 37⁰C. The next day, cells were treated with 100 µg/mL concentration of cycloheximide (CHX) (Sigma-Aldrich, US) or with fresh media. The plate was incubated at 37⁰C for 1h and 5min to allow for activation of the inhibitor. Cells were treated with TRI reagent (0.3-0.4mL per 1x10^5^-10^7^ cells) from Invitrogen (Carsbad, CA) and RNA extraction performed according to manufacturer’s instructions. Quantitative reverse transcription PCR (RT-qPCR) was performed as described.[24] Endothelial expression of exons was also examined in whole transcriptome data from BOECs, following conversion of genomic coordinates from GRCh18 to GRCh38.[37] Binary sequence alignment map (bam) files aligned to GRCh38[37] were analyzed in Galaxy Version 2.4.1[38] and the Integrated Genome Browser (IGB) 9.1.8.[39]

To examine broader relevance, selected genes were examined for bulk RNASeq expression across the 53 tissues in the Genotype-Tissue Expression (GTEx) project,[40] supplemented by expression in normal human peripheral blood mononuclear cells (PBMCs) treated with 100µg/mL CHX for 1h as described elsewhere.[41]

### Statistical analysis

Creation of customised scripts and data analysis was performed in R. Subsidiary analyses and generation of graphics used GraphPad Prism 9 (GraphPad Software, San Diego, CA). Two group comparisons were performed using Mann Whitney rank sum. Three group comparisons were performed using Kruskal Wallis, with Dunn’s post test applied for selected pairwise comparisons.

## RESULTS

### Endothelial cell morphology after 1h and 3h cycloheximide

In three separate culture pairs of HMECs pre and post 1h (1 pair) or 3h (2 pairs) CHX treatment, no clear morphological differences were apparent (Figure S1). In contrast HPAECs displayed no morphological differences after 1h CHX but some minor morphological changes after 3h (Figure S1). We concluded that under the culture conditions used, HMEC were a more resilient primary endothelial cell type than HPAEC, and potentially suitable for 1h and 3h treatments. 1h was selected to better facilitate capture of acute responses to short term inhibition of NMD and translation, while minimising potential confounding of secondary consequences due to later initiation of secondary and apoptotic pathways.

### 1h cycloheximide-differentially expressed genes replicated by longer, NMD-specific depletion strategies in other cell types

In the HMECs, RNAseq identified alignments to multiple RNA types, with a similar distribution in cycloheximide and media-treated endothelial cells. 419 genes displayed differential alignments to p<0.15 after 1h cycloheximide treatment (Figure 1Bi). Of these, 28 (6.7%) were in the 1,493 genes differentially expressed >12 days after *SMG5*, *SMG6* or *SMG7* knockdown in mouse embryonic stem cells (mESCs, Table S1A).[34] Similarly, of the current study’s 419 most differentially expressed genes, 9 (2.2%) were in the 357 genes differentially expressed 72hs after siRNA inhibition of *UPF1* in HeLa cells[35] (Figure 1Bi, Table S1B). These 9 included *PDE4B* which was also differentially expressed in [34], and *MAP3K14*, one of the two genes where Mendell et al[35] demonstrated the relative increase in half-life to be >7-fold longer after UPF1 inhibition. Overall, the current dataset provided a similar proportion of overlapping genes to the mESCs[34] as the 72h HeLa study[35] despite a considerably shorter time period (1h) and less specific mechanism of NMD inhibition (Figure 1Bii).

### Biological processes identified by analysis of differentially expressed genes after 1h cycloheximide

For the current 1h cycloheximide-treated HMEC dataset, David GO term clustering of genes differentially expressed to p<0.15 after cycloheximide identified biological processes (Table S1C). The highest scoring clusters were found to be involved in cell adhesion, response to lipopolysaccharides, amino acid transport, and the integrator complex (Table S1C). In contrast, clustering of the same number of random genes did not identify any significant processes.

To determine if the specificity could be enhanced, processes identified by genes differentially expressed in HDMEC and also identified in earlier studies inhibiting NMD were examined (Table 1). Despite the small number of genes (N=36), a significant cluster was generated, comprising genes mapping to terms related to the C-terminal domain found in proteins belonging to two of the five helicase superfamilies.

**Table 1.** Representative terms significantly clustered from 37 genes differentially expressed in multiple NMD datasets.

| Source | Term | Genes | p-value | Benjamini p-value |
| --- | --- | --- | --- | --- |
| UP_SEQ_FEATURE DOMAIN: | Helicase C-terminal | *DHX57, CHD6, INO80, SMARCA1* | 7.640E-4 | 0.078 |
| INTERPRO, IPR014001 | Helicase, superfamily 1/2, ATP-binding domain 4 | *DHX57, CHD6, INO80, SMARCA1* | 9.959E-4 | 0.062 |
| SMART, SM00490 | Helicase superfamily c-terminal domain, HELICc 4 | *DHX57, CHD6, INO80, SMARCA1* | 0.00137 | 0.023 |
| GOTERM_MF_DIRECT GO:0008094 | DNA-dependent ATPase activity | *CHD6, INO80, SMARCA1* | 0.0020 | 0.21 |
| GOTERM_MF_DIRECT GO:0005524 | ATP binding 9 | *DHX57, ADK, CHD6, ABCA7, INO80, SMARCA1, ABCC10, CLK4, MAP3K14* | 0.0034 | 0.21 |
| INTERPRO, IPR027417 | P-loop containing nucleoside triphosphate hydrolase | *DHX57, CHD6, ABCA7, INO80, SMARCA1, ABCC10, RHOBTB1* | 0.0051 | 0.12 |
| UP_KW_LIGAND KW-0067 | ATP-binding | *DHX57, ADK, CHD6, ABCA7, INO80, SMARCA1, ABCC10, CLK4, MAP3K14* | 0.0058 | 0.03 |

### Cellular processes targeted by miRNAs

Next, we examined RNAseq alignments to pre-miRNAs in the primary human endothelial cells (HMEC). There were alignments to 113 pre-miRNAs, and of these, 34/113 (30%) displayed expression levels that differed by at least 20% between media-and cycloheximide-treated cells. Thirteen pre-miRNAs were higher in media treated cells, and 21 in cycloheximide-treated cells. Target genes of these pre-miRNAs were identified, taking into consideration that pre-miRNAs can give rise to two mature forms of miRNAs (5’ and 3’), in which case, targets of both miRNAs were determined. GO term clustering was performed on these genes and the most frequently targeted processes are shown in Figure 2A. Notably, nearly one-third (31.6%) of miRNAs target genes were found to be involved in the regulation of phosphorylation, one-sixth (13%) in phosphatidylinositol signalling, one-sixth (13%) in transcription and one-twentieth (5.2%) in cell adhesion, mirroring processes identified by mRNAs.

**Figure 2:**
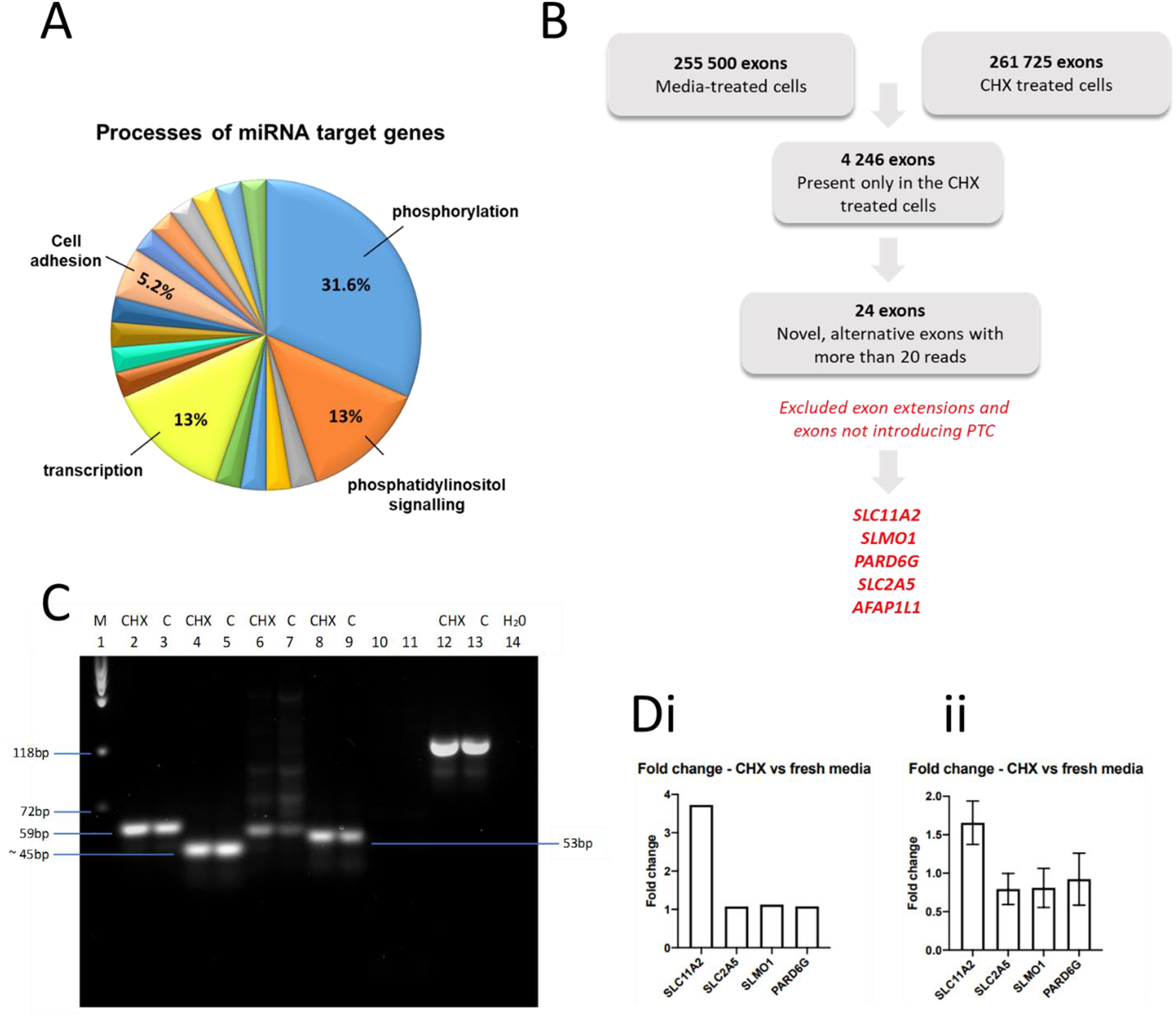
qPCR validations of endothelial cell NMD. **A. Gene ontology processes implicated by pre-miRNAs differentially expressed in cycloheximide-treated endothelial cells.** Gene ontology (GO) term clustering was performed on the mRNA targets of 34 miRNAs whose expression differed by at least 20% between cycloheximide and media treated HMEC cells. **B. Selection of exons likely to be targeted by NMD.** Data from media-treated and cycloheximide-treated endothelial cells were compared to identify exons where alignments were only identified in cycloheximide-treated cells and were novel, alternative exons that introduce a PTC into the transcript. Sequences and chromosomal position for all 24 novel alternative exons are provided in Table S2**. C. Validation methodology** by low melting agarose gel electrophoresis of DNA fragments from HUVECs treated with cycloheximide or fresh media, amplified by RT-PCR using primers designed for exons likely targeted by nonsense mediated decay (NMD). Lanes: 1, ΦX174 DNA marker; 2,3, FSLC11A2 and RSLC11A2; 4,5, FSLC2A5 and RSLC2A5; 6,7, FSLMO1 and RSLMO1; 8,9, FPARD6G and RPARD6G; 12,13, GAPDH; 14, water control. Abbreviations: CHX, cycloheximide, C, control (fresh media), M, DNA marker. **D. Validations** by fold change in expression of exons in cycloheximide-treated EC. **i)** HUVEC and **ii)** BOEC treated with 100µg/ml cycloheximide or fresh media for 1h, and expression of candidate exons compared, normalised to reference gene *GAPDH*. Data from *AFAP1L* are not shown as this demonstrated low level of amplification using all tested primers and ultimately it was not included in the final analysis.

### Exons with multiple alignments only in endothelial cells treated with 1h cycloheximide

To identify specific exons that may be targeted experimentally as assays of active NMD/translation inhibition in qRT-PCR or short-read RNASeq analyses, individual exon alignments were examined in the endothelial RNAseq datasets. There were 255,500 alignments to exons in the media-treated cells and 261,725 alignments in cycloheximide-treated cells. Comparison of exon alignments identified 4,246 exons observed only in the cycloheximide-treated cells (Figure 2B). After applying the criteria for selection of novel, alternative exons with more than 20 reads, the list was reduced to 24 exons (Table S2). Following this, exons of genes with poorly characterised functions, exon extensions and exons that neither had a stop codon in each reading frame nor induced frameshift leading to the formation of a premature stop codon in the downstream exon were excluded. This allowed the construction of a final prioritized list that comprised 5 exons from *SLC11A2, SLC2A5, SLMO1, PARD6G* and *AFAP1L*.

### Validation of HMEC exons in other endothelial cell types

To test if findings were specific to human microvascular endothelial cells (HMEC), qRT-PCR validations were performed on cDNA from human umbilical vein endothelial cells (HUVEC) and blood outgrowth endothelial cells (BOECs) following treatment with cycloheximide or fresh media for 1h. Oligonucleotide primers were designed for the alternative exons inducing PTC formation in *SLC11A, SLC2A5, SLMO1*, *PARD6G* and *AFAP1L1*. GAPDH allowed for relative quantification of expression in both assessed conditions. qRT-PCR products were size-separated using low melt agarose gel electrophoresis to confirm nucleic acid sizes (Figure 2C): Amplicons produced by *FSLC11A2* and *RSLC11A2, FSLMO1* and *RSLMO1,* and *FPARD6G* and *RPARD6G* were consistent with the expected sizes although *FSLC2A5* and *RSLC2A5* designed to target the alternative U8 exon in the *SLC2A5* gene initially amplified a smaller fragment than expected. The ΔΔCt method was used to determine the fold change in expression of single band exons of interest in cycloheximide-treated HUVEC (Figure 2Di) and BOECs (Figure 2Dii). In crude and GAPDH-normalized analyses, expression of the 3B exon from *SLC11A2* emerged as consistently higher in all endothelial cell types examined after treatment with cycloheximide, at almost 4-fold higher in cycloheximide-treated HUVEC (Figure 2Di) and with a 1.7-fold increase in BOECs (Figure 2Dii). Analyses of the remaining exons did not indicate any consistent changes across all endothelial cell types (Figure 2D).

### Validation of HMEC exons in primary blood outgrowth endothelial cells (BOECs) treated with cycloheximide

If differentially-expressed exons are NMD targets, then the RNASeq alignments should be to fully spliced exons.[1–3] Binary alignment map (bam) files demonstrated that for *SLC11A2, PARD6G* and *SLMO1*, the exon sequences were expressed in a small proportion of human primary BOECs derived from 4 different donors (Figure 3A). *SLC11A2* exon 3B was expressed consistently, at a low level compared to the adjacent exon (mean 5.9%, SD 2.0%, Figure 3B). Examining exact bam file alignments, only *SLC11A2* exon 3B exhibited exact alignments indicative of previous splicing, and this pattern was observed in BOECs from 3 separate donors (Figure 3C).

**Figure 3:**
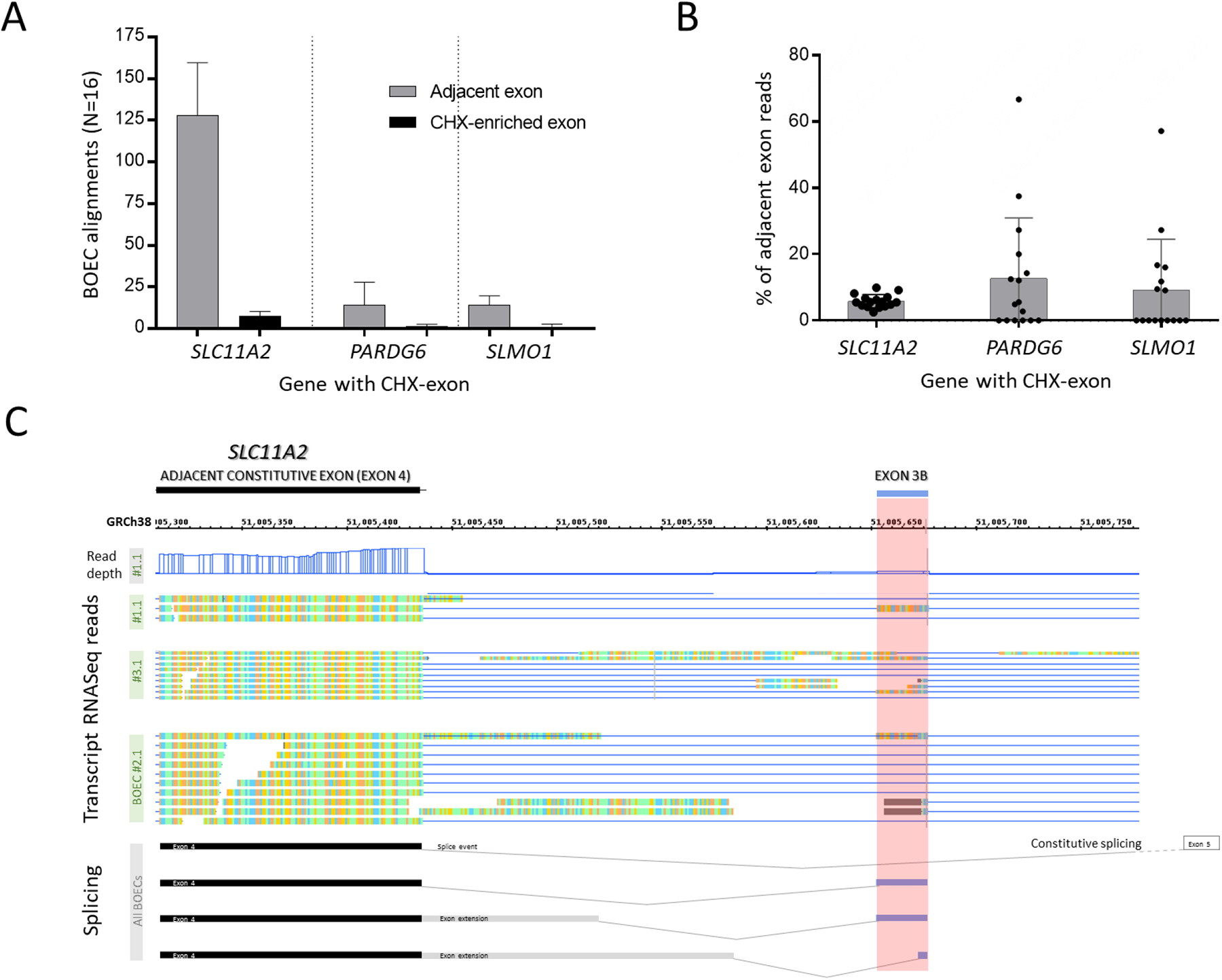
RNASeq validation of NMD-target exons in blood outgrowth endothelial cells. **A)** Alignments to the cycloheximide-enriched exon, and nearest adjacent exon for 3 separate genes, across 16 BOECs from 4 separate donors, as described in [36]. Mean and standard deviation displayed for **i)** absolute alignments and **ii)** percentage of alignments to the nearest adjacent exon. **B)** Integrated Genome Browser (IGB) 9.1.8 views of GRCh38 chr12:51,005,295-51,005,797 spanning the cycloheximide enriched exon 3B and adjacent constitutive exon 4 of *SLC11A2*. The position of exon 3B is highlighted in red across GRCh38 coordinates, 3 separate BOEC sets of alignments and transcript cartoons. Note the sharp boundaries of alignments to exon3B, and its absence from the RefSeq curated transcript denoted by constitutive splicing cartoon.

Three other alignment patterns were observed in the BOECs in addition to the precise splicing (pattern 1) seen for *SLC11A2* exon 3B: 5’ extension (pattern 2), both 5’ and 3’ extension (Pattern 3), and use as an alternate first exon (Pattern 4, Figure 4). *SLC11A2* exon 3B displayed predominantly patterns 1 and 2, *PARD6G* exon 2A predominantly Pattern 3, and *SLM01* exon 1B predominantly Pattern 4. We concluded that the novel *SLC11A2* exon 3B was in a transcript most consistently stabilized by NMD inhibition.

**Figure 4:**
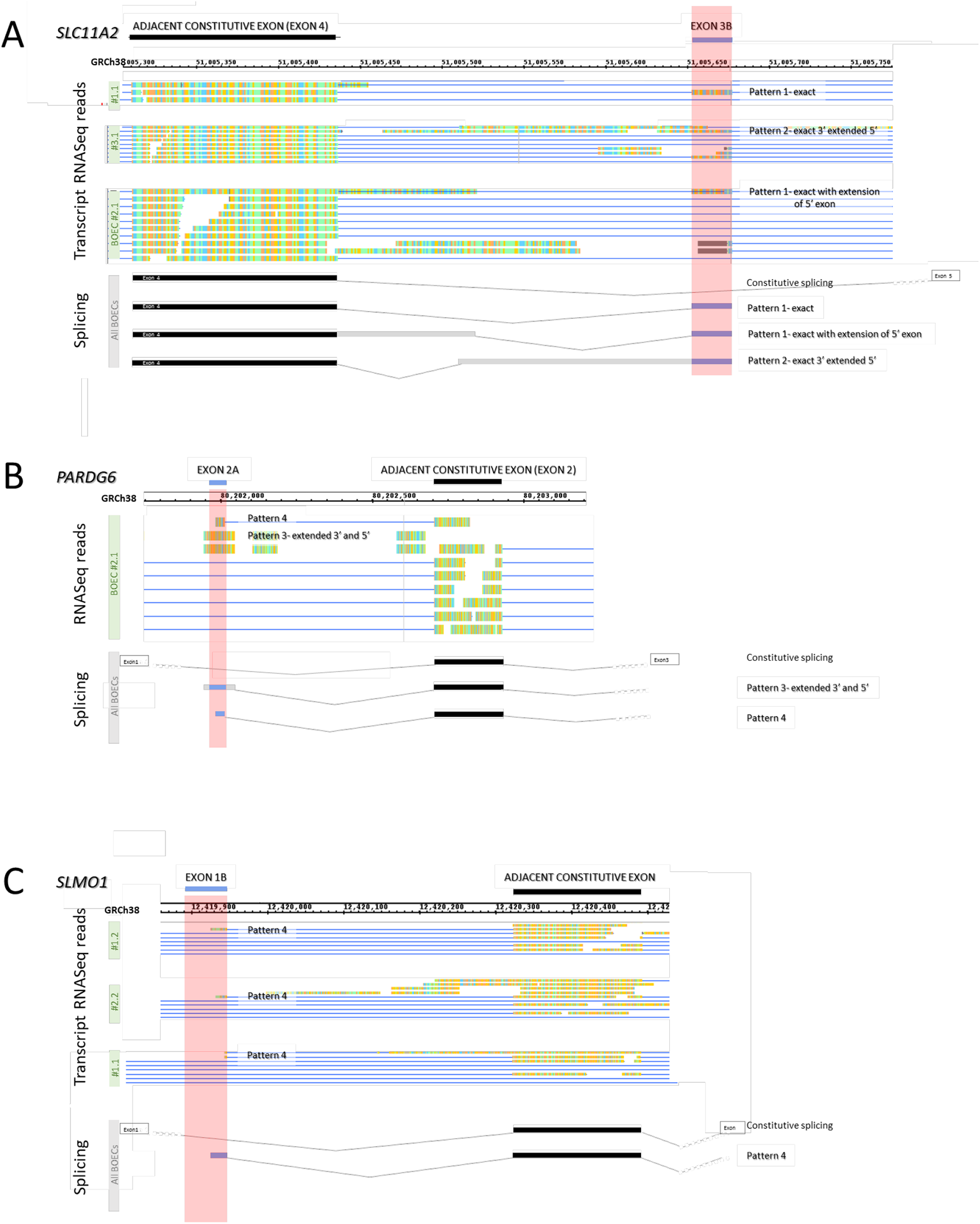
Patterns of RNASeq alignments to NMD-target exons in blood outgrowth endothelial cells Integrated Genome Browser (IGB) 9.1.8 views of GRCh38/hg38 and transcript cartoons. A) chr12:51,005,295-51,005,797 spanning the cycloheximide (CHX)-enriched exon 3B and adjacent constitutive exon 4 of *SLC11A2* (same reads as in Fig. 5C, now annotated by pattern type; B) chr18:80,200,939-80,203,651 spanning the CHX-enriched exon 2A and adjacent constitutive exon 2 of *PARD6G*; C) chr18:12,419,463-12,420,541 spanning the CHX-enriched exon 1B and adjacent constitutive exon of *SLMO1*. For each gene, the position of the CHX-enriched exons are highlighted in red across separate BOEC sets of alignments for A) *SLC11A2* exon 3B; B) *PARD6G* exon 2A; C) *SLMO1* exon 1B. Note exon-exon boundary read through in B) and C), and CHX-enriched exons absent from respective RefSeq transcripts denoted by constitutive splicing patterns.

### Broader relevance of *SCL11A2*

*SLC11A2* is expressed at modest levels (median 4.05-35.21 transcripts per million (TPM)) in bulk RNASeq analyses across 53 tissues in the Genotype-Tissue Expression Project (GTEx).[40] *SLC11A2* (also known as *NRAMP2* or *DMT1*) is recognised as encoding an essential iron transporter where common[42] and rare[43] variants result in microcytic anaemia with iron overload. More recently, *SLCA11A2* overexpression has been reported as a promising biomarker and therapeutic target in ovarian cancer.[44] While alternate 5’ and 3’ splicing is known to critically regulate *SLCA11A2* subcellular location and site of iron transport,[45,46] alternate splicing is more extensive. In the course of this project, exon 3B was identified within a RefSeq non-coding RNA transcript, NR_183176.1 [*SLC11A2* transcript variant 25] which was originally identified in a Roswell Park Cancer Institute Human BAC Library.[47] To further support wider roles, in separate rRNA-depleted RNASeq libraries from peripheral blood mononuclear cells resuspended in endogenous plasma with and without 1h cycloheximide 100μg/ml,[41] *SLC11A2* was differentially expressed after cycloheximide (Supplementary Figure 2A). Low level expression of exon 3B sequences were identified, though there were no significant changes by DEXSeq after 1h cycloheximide (Supplementary Figure 2B). Further studies are ongoing to evaluate the effects of short term cycloheximide in this cell type and experimental system (Li et al, manuscript in preparation).

## DISCUSSION

Understanding when NMD and translation inhibition is operational is important for assessment of general cellular responses, in addition to evaluating impacts from NMD-based therapeutics. Here we have added to existing data regarding genes and biological processes likely to be impacted by NMD inhibition through analysis of differentially-expressed genes, and target mRNAs of differentially-expressed miRNAs, following 1h cycloheximide treatment of a series of different endothelial cell types. The study also identified specific exons likely targeted by NMD, confirmed relevance in a number of additional cell types, and provided cell biologists with a laboratory validation assay for HUVEC and BOECs.

A potential weakness of the study was the use of cycloheximide which can cause cell death in prolonged treatment. This was mitigated during experimental design to select the most resilient endothelial cell type, and by restricting cycloheximide treatment to 1h (Figure S1). Future studies may examine endothelial cells’ acute responses by inhibiting NMD by other methods such as siRNA or CRISPR/Cas9 to knock down individual NMD factors. That said, strengths of the current study included overlap with 37 genes shown to be differentially expressed in other cell types and species after targeting individual NMD factors (Figure 1B, Table S1). Huth et al[34] knocked down SMG5, SMG6, and SMG7 in mouse embryonic stem cells (ESCs) and analysed after 12 and 15 days,[48] while Mendell et al depleted UPF1 in human HeLa cells and analysed after 72h.[35] The paucity of shared genes differentially expressed by both of these datasets has been attributed to different cell types and NMD inhibition methods. The current dataset provided a similar proportion of overlapping genes (Figure 1, Table S1), despite previous datasets representing pluripotential cells,[34] and an immortalized cell line established in 1951 that does not have a normal karyotype.[35] We speculate that the current study may present results closer to real biological differences for differentiated human cells responding to physiological inhibitions of NMD because of using primary human microvascular endothelial cells. Further, previous studies examined cells after many days of NMD inhibition whereas the current study restricted to 1h, a state more analogous to the suppression of protein translation and NMD following phosphorylation of the α subunit of eukaryotic initiation factor 2 (eIF2α) as part of the integrated stress response that aims to restore cellular homeostasis following a variety of cellular stresses.[49–52]

Further study strengths include identification of processes previously described to be associated with NMD. Identification of the integrator complex is highly relevant to recent data indicating that the integrator complex regulates production of full length coding and non-coding RNAs by mechanisms including RNA cleavage,[53] RNA polymerase II pausing,[54] premature transcriptional termination,[55] and RISC loading of miRNAs.[56] ATP is relevant particularly to UPF1 which has both ATPase and RNA helicase activities.[7] Additional biological functions were elucidated by examining the targets of miRNAs differentially expressed in cycloheximide and media treated cells – phosphorylation, phosphatidylinositol signalling, transcription, and cell adhesion were in-keeping with more recent data in other cell types that is leading to more targeted therapeutics.[57–59] Again, this was further supported by clustering the genes that were also differentially expressed in mouse ESCs[34] or HeLa cells[35] after different methods of NMD inhibition (Table 1).

A specific strength of this study is the identification of endothelial-expressed exons likely stabilised by NMD inhibition, determined using very stringent selection criteria that reduced the number of candidate exons from 4,246 to 24 (Table S2) then only 5 before validation in HUVEC and BOECs. For *SLC11A2*, there was a difference between the level of exon 3B expression in media-treated HMECs (absent), HUVECs/ BOECs where designed primers amplified their target sequences, and PBMCs where expression was at a higher level. Potential reasons for differences may relate to the exon’s inclusion in a transcript not subject to NMD in other cell types, or varying activity of NMD at the time of the experiment, for example reflecting differing regulation of NMD in mammalian cell lineages[60,61], or prevailing stress,[9,10] and we speculate, differing proportion of serum in culture conditions.

For specific exons, the 3B alternative exon in *SLC11A2* was not detected at baseline in HMEC, was in a transcript stabilised by cycloheximide treatment of HMEC and increased almost 4-fold in cycloheximide-treated HUVEC. Further, it displayed precise and consistent alignments in BOECs from separate donors, is now known to be part of RefSeq non-coding RNA transcript NR_183176.1, and displays higher proportional expression in peripheral blood mononuclear cells (PBMCs). Since *SLC11A2* encodes a divalent metal transporter 1 (DMT1) responsible for absorption and transport of metals including cadmium, copper, zinc, iron and manganese, inclusion in toxicity assays based on these treatments[26] may prove valuable, in addition to the wider proposed use as positive controls for experiments assessing the efficiency of NMD-targeting compounds in patient-derived endothelial cells.

In conclusion, the study contributes to our understanding of natural targets regulated by nonsense mediated decay and protein translation inhibition. Interactions of NMD with cellular pathways were identified through analysis of differentially expressed genes, and target genes of miRNAs exhibiting differential expression. The data highlight the variability of activity in different tissues and cells, and the datasets and exons provide useful tools for positive controls for future studies assessing efficiency and toxicology of NMD-targeting compounds in patient-derived cells.

## Acknowledgments

A.M.B received specific fellowship funding from the Association of Physicians of Great Britain and Northern Ireland. The microvascular endothelial cell RNASeq libraries were established with funding support from the British Heart Foundation (PG/09/041/27515). Blood outgrowth endothelial cells (BOECs) were established with funding support from the National Institute for Health Research Imperial Biomedical Research Centre, London, UK, and earlier, from the Congressionally Directed Medical Research Programme of the United States Department of Defense (W81XWH-16-1-0607). BOEC RNA Sequencing was performed with funding from the Imperial College Healthcare NHS Trust. Peripheral blood mononuclear cells were established and sequenced with funding from the D’Almeida Charitable Trust, UK.

## Author contributions

Conceptualization, A.M.B., F.S.G., I.G.M. and C.L.S.; methodology, A.M.B., F.S.G., D.P., M.E.B-H., M.A.A., I.G.M. and C.L.S.; software, I.G.M.; validation, A.M.B., D.P., D.L. and C.L.S.; formal analysis, A.M.B., I.G.M. and C.L.S.; investigation, A.M.B., F.S.G., D.P., M.E.B-H., D.L., M.A.A., I.G.M. and C.L.S.; data curation, I.G.M. and C.L.S.; writing—original draft preparation, A.M.B.; writing—review and editing, A.M.B.; M.E.B-H..; I.G.M.; C.L.S.; visualization, A.M.B. (Figures 1A, 2; Tables S1, S2) and C.L.S. (Figures 1B, 3, 4; Tables 1, S1); supervision, I.G.M.; C.L.S.; project administration, I.G.M. and C.L.S.; funding acquisition, A.M.B., M.A.A. and C.L.S. All authors have read and agreed to the published version of the manuscript.

## Data availability

Non-sensitive data underlying this article are to be made available at Zenodo to be used under the Creative Commons Attribution licence. Primary sequence data used in this research was collected subject to the participants’ informed consent. Access to these data will only be granted in line with that consent, subject to approval by the project ethics board and under a Data Sharing Agreement. Blood outgrowth endothelial cells used in this research were collected subject to the informed consent of the participants. Access will only be granted in line with that consent, subject to approval by the project ethics board.

