## Supplementary Data for "Transcriptome-wide analysis of primary human endothelial cell responses to 1 hour of protein translation inhibition identify nonsense mediated decay targets and a non-coding *SLC11A2* exon as an acute biomarker"

### DATA SUPPLEMENT:

#### SUPPLEMENTARY FIGURES

**Figure S1:** Morphologial appearances of endothelial cells treated with fresh media or media supplemented with 100µg/mL cycloheximide for 1h or 3h.

**Figure S2:** *SLC11A2* expression and splicing in human peripheral blood mononuclear cells compared to endothelial cells.

#### SUPPLEMENTARY TABLES

**Table S1:** Genes and processes identified in studies inhibiting nonsense mediated decay

**Table S2:** Novel alternate exons in cycloheximide-treated HMEC

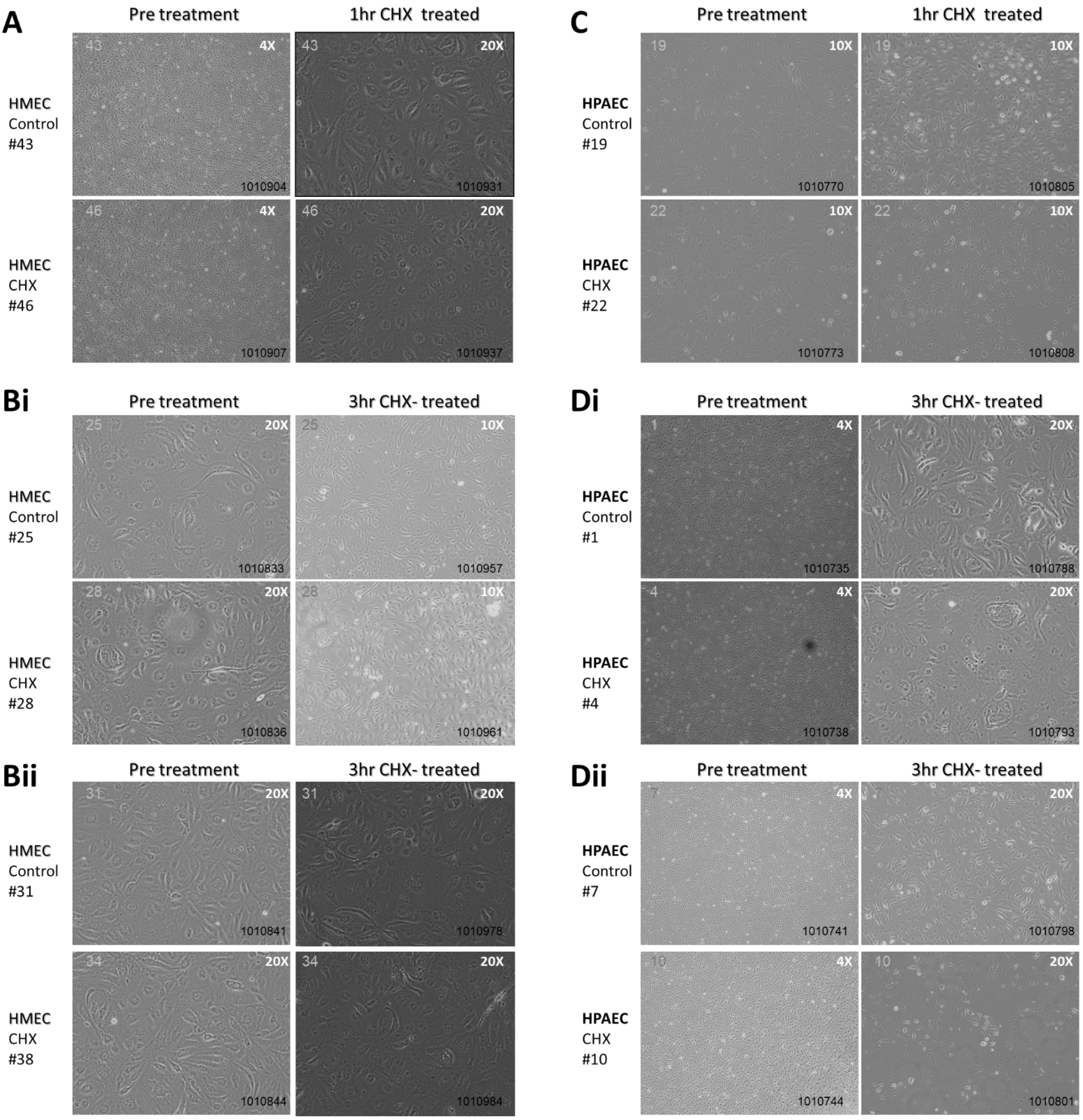


**Figure S1: Morphologial appearances of endothelial cells treated with fresh media or media supplemented with 100ug/mL cycloheximide for 1h or 3h.**

To optimise treatment times, during experimental design, primary human microvascular EC (HMEC) and primary human pulmonary artery EC (HPAEC) from Promocell (PromoCell GmbH, Heidelberg) were cultured in the presence and absence of cycloheximide (CHX) 100µg/mL for 1h or 3h, and photographed prior to harvesting in Tri reagent (Cambridge Bioscience Ltd, Cambridge, UK) and immediate storage at -70ºC**.** **A)** HMEC pre and post 1h treatment –no morphological differences were apparent. **B)** Two separate culture pairs of HMEC cultures pre and post 3h treatment – again no morphological differences were apparent. **C)** HPAEC pre and post 1h treatment. The comment recorded on the day of treatment indicated no morphological differences were apparent. **D)** Two separate culture pairs of HPAEC cultures pre and post 3h treatment. Comments recorded on day of treatment for both **i)** 1010793, and **ii)** 1010801, were of morphological changes, and significant cell detachment is evident.

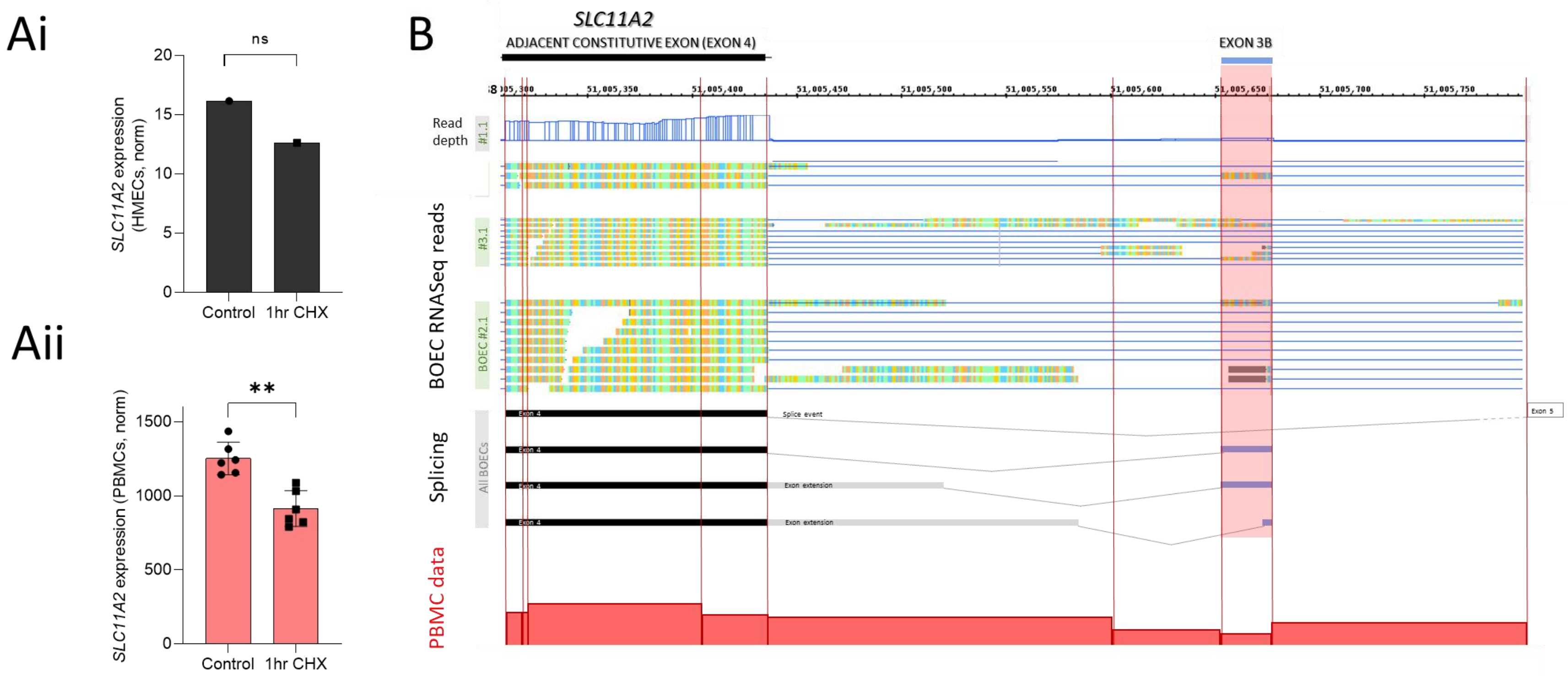


**Figure S2: *SLC11A2* expressions patterns in human peripheral blood mononuclear cells (PBMCs) compared to endothelial cells.** As described by Xiao et al,[1] freshly derived PBMCs were obtained from 6 donors, resuspended in endogenous plasma, and distributed to separate experimental treatment tubes each containing ~5ml of cell/plasma suspension. For each donor, one set of tubes were treated with <100µl cycloheximide (CHX) to a final concentration of 100µg/ml, and a second set with <100µl of Dulbeccos Modified Eagle Medium (DMEM) supplemented with 10% fetal calf serum (FCS). PBMCs were cultured at 37ºC for 1hr before centrifugation, resuspension of the cell pellet in Tri reagent, RNA extraction, ribosomal (r)RNA depletion, and generation of libraries for Illumina HiSeq sequencing using paired-end 150bp reads (Genewiz, Leipzig, Germany). Genewiz performed differential gene expression analyses using DESeq2,[2] and differential exon expression using DEXSeq[3] on exon regions and junctions of all genes. **A) Overall expression of *SLC11A2*. i) Endothelial:** Mean alignments in the original endothelial experiment in HMEC, quantified across 18 exons and junctions in HMEC treated with fresh media for 1h, and 20 exons and junctions in HMEC after 1h treatment with 100µg/ml cycloheximide (CHX). The trend for fewer alignments post CHX did not meet significance (p=0.51). **ii) PBMC**: DESeq normalised mean reads for *SLC11A2* expression in PBMCs resuspended in endogenous plasma after 1h incubation with and without 100µg/ml cycloheximide (CHX). Note the similar decrement post CHX and that in this cell type, using mean data from replicates in 6 donors, the p-value calculated by Mann Whitney was 0.0022 (**). For the other HMEC genes where differentially-spliced exons were identified post cycloheximide, *PARD6G* and *SLMO1* (*PRELID3B*) were expresssed in PBMCs but displayed no differential expression after 1h cycloheximide. **B) Alignments across GRCh38 chr12:51,005,295-51,005,797 spanning the cycloheximide-enriched exon 3B and adjacent constitutive exon 4 of *SLC11A2*.** Upper panels replicate data shown in Figure 2D, lowest panel (red bars), the mean expression for bases per exon region in PBMCs from 3 donors calculated by DEXSeq,[3] with exon regions demarcated by red vertical lines. None of this region met DEXSeq thresholds for differential expression after 1h cycloheximide in PBMCs, although reduced alignments post CHX were observed elsewhere in the gene.

*[1] Xiao S, Kai Z, Murphy D, Li D, Murphy D, Li D, et al. Am J Hum Genet 2023 In Press*
*[2] Anders S, Huber W. Genome Biol 2010, 11(10):R106*
*[3] Anders S, Reyes A, Huber W. Genome Res 2012, 22(10):2008—17*

**Table S1: Genes and processes identified in studies inhibiting nonsense mediated decay:**

| **GENES SHARED** | **Chr.** | **Gene description** | **HMEC data** | | | |
|---|---|---|---|---|---|---|
| **A) Mouse cells** [5] | | | **Mean control** | **Mean at 1h CHX** | **Fold change** | **P value** |
| *ABCA7* | chr19 | ATP-binding cassette, sub-family A, member 7 | 1.86 | 4.41 | 2.37 | 0.012 |
| *ABCC10* | chr6 | ATP-binding cassette, sub-family C, member 10 | 2.44 | 4.42 | 1.81 | 0.083 |
| *ADK* | chr10 | adenosine kinase isoform b | 8.51 | 16.25 | 1.91 | 0.097 |
| *BMP4* | chr14 | bone morphogenetic protein 4 preproprotein | 3.96 | 7.24 | 1.83 | 0.089 |
| *CCDC134* | chr22 | coiled-coil domain containing 134 | 5.65 | 2.57 | 0.46 | 0.069 |
| *CHD6* | chr20 | chromodomain helicase DNA binding protein 6 | 7.17 | 9.36 | 1.31 | 0.076 |
| *CLK4* | chr5 | CDC-like kinase 4 | 5.05 | 8.42 | 1.67 | 0.099 |
| *DHX57* | chr2 | DEAH (Asp-Glu-Ala-Asp/His) box polypeptide 57 | 6.24 | 4.67 | 0.75 | 0.072 |
| *DOCK6* | chr19 | dedicator of cytokinesis 6 | 12.13 | 18.92 | 1.56 | 0.059 |
| *INO80* | chr15 | INO80 complex homolog 1 | 3.60 | 5.83 | 1.62 | 0.097 |
| *LRIG2* | chr1 | leucine-rich repeats and immunoglobulin-like | 5.89 | 10.51 | 1.79 | 0.083 |
| *LTBP3* | chr11 | latent transforming growth factor beta binding | 7.20 | 4.02 | 0.56 | 0.027 |
| *MPHOSPH9* | chr12 | M-phase phosphoprotein 9 | 5.86 | 9.77 | 1.67 | 0.080 |
| ***PDE4B*** | chr1 | phosphodiesterase 4B, cAMP-specific isoform 2 | 8.30 | 25.04 | 3.02 | 0.002 |
| *PRKAB2* | chr1 | AMP-activated protein kinase beta 2 | 4.41 | 2.79 | 0.63 | 0.078 |
| *RHBDD2* | chr7 | rhomboid domain containing 2 isoform b | 3.10 | 4.97 | 1.60 | 0.018 |
| *RHOBTB1* | chr10 | Rho-related BTB domain containing 1 | 11.33 | 6.52 | 0.58 | 0.039 |
| *SH3TC1* | chr4 | SH3 domain and tetratricopeptide repeats 1 | 11.19 | 16.92 | 1.51 | 0.062 |
| *SMARCA1* | chrX | SWI/SNF-related matrix-associated | 9.94 | 14.23 | 1.43 | 0.073 |
| *SPIRE2* | chr16 | spire homolog 2 | 23.76 | 12.32 | 0.52 | 0.082 |
| *STX1A* | chr7 | syntaxin 1A (brain) | 18.91 | 30.85 | 1.63 | 0.075 |
| *TATDN1* | chr8 | TatD DNase domain containing 1 | 5.76 | 9.04 | 1.57 | 0.087 |
| *TMEM63C* | chr14 | transmembrane protein 63C | 4.02 | 2.44 | 0.61 | 0.145 |
| *TTC14* | chr3 | tetratricopeptide repeat domain 14 isoform b | 17.92 | 10.01 | 0.56 | 0.048 |
| *TTC39C* | chr18 | tetratricopeptide repeat domain 39C | 1.65 | 8.09 | 4.92 | 0.018 |
| *ZC3H12A* | chr1 | zinc finger CCCH-type containing 12A | 3.52 | 6.09 | 1.73 | 0.042 |
| *ZFAND2A* | chr7 | zinc finger, AN1-type domain 2A | 15.62 | 6.66* | 0.43 | 0.122 |
| *ZZEF1* | chr17 | zinc finger, ZZ type with EF hand domain 1 | 8.24 | 10.96 | 1.33 | 0.090 |
| **B) HeLa Cells** [7] | | | | | | |
| *ADM* | chr11 | adrenomedullin | 6.30 | 14.20 | 2.26 | 0.083 |
| *AGL* | chr1 | amylo-1, 6-glucosidase, | 4.17 | 5.47 | 1.31 | 0.076 |
| *FLNA* | chrX | filamin A, alpha isoform 1 | 44.21 | 55.85 | 1.26 | 0.098 |
| *HDAC3* | chr5 | histone deacetylase 3 | 16.14 | 8.10 | 0.50 | 0.087 |
| *MAP3K14* | chr17 | mitogen-activated protein kinase kinase kinase | 1.40 | 3.97 | 2.84 | 0.075 |
| ***PDE4B*** | chr1 | phosphodiesterase 4B, cAMP-specific isoform 2 | 8.30 | 25.04 | 3.02 | 0.002 |
| *SLC36A1* | chr5 | solute carrier family 36 member 1 | 3.10 | 5.83 | 1.88 | 0.136 |
| *SLC9A3R1* | chr17 | solute carrier family 9 Subfamily A (NHERF1) | 3.18 | 9.59 | 3.02 | 0.099 |
| *THOC1* | chr18 | THO complex 1 | 7.09 | 10.50 | 1.48 | 0.090 |

| **C) GO PROCESSES (HMEC only)** | **Highest scoring GO term for annotation clusters** | **Enrichment Score** | **Term p-value** |
|---|---|---|---|
| ***Cluster and term met predesignated significance thresholds:*** | | | |
| GO:0007156~homophilic cell adhesion via plasma membrane adhesion molecules | | 4.34 | 1.37E-07 |
| GO:0071222~cellular response to lipopolysaccharide | | 1.96 | 9.95E-05 |
| GO:0006865~amino acid transport | | 1.32 | 0.033 |
| GO:0032039~integrator complex | | 1.18 | 0.025 |
| ***Cluster and term did not meet predesignated significance thresholds:*** | | | |
| *GO:0032508~DNA duplex unwinding* | | *0.90* | *0.059* |
| *GO:0090263~positive regulation of canonical Wnt signaling pathway* | | *0.62* | *0.097* |
| *GO:0001934~positive regulation of protein phosphorylation* | | *0.50* | *0.115* |
| *GO:0008380~RNA splicing* | | *0.38* | *0.121* |
| *GO:0007264~small GTPase mediated signal transduction* | | *0.42* | *0.127* |

**Legend:** Expression and gene ontology (GO) data for genes identified as differentially expressed to p<0.15 in primary human microvascular endothelial cells (HMEC), indicating mean reads to exons and exon junctions in control HMEC, and HMEC treated with cycloheximide (CHX) for 1hr; fold difference, and p value calculated by Welch t-test as in [4]. **A)** Genes that were also differentially expressed in Huth et al [5] following knockdown of NMD factors SMG5, SMG6 or SMG7 in mouse embryonic stem cells (mESCs) cultured for >12 days, as described.[6] **B)** Genes that were also differentially expressed in Mendell et al, examining *UPF*-depeleted HeLa cells.[7] **C)** Highest scoring Gene Ontology (GO) terms.[8,9] Processes from genes also in mESC or HeLa lists are provided in main Table 1.

### Table S2: Novel alternate exons in cycloheximide-treated HMEC

| Gene | ExonID | GeneID | Exon | Length | Chr | Strand | Start | End | Sequence |
|---|---|---|---|---|---|---|---|---|---|
| *SLC11A2* | 199534 | 12155 | 3B | 25 | chr12 | - | 49685703 | 49685727 | AATAAGAGGCTGATGGAACCTGCAG |
| *SLC2A5* | 26066 | 1636 | U8 | 56 | chr1 | - | 9054950 | 9055005 | GCTGGAGTGCAGTGGCATGATCGAATTCCTGGGCTCAAGCGATCCTCTTGCCTCAG |
| *PARDG6* | 262884 | 15890 | 2A | 55 | chr18 | - | 76060838 | 76060892 | GTGGTGAACACTGACCACCTCAAGGAACCAAGCAGCCAGTGGTGACAGTTGCAAG |
| *SLMO1* | 263721 | 15939 | 1B | 56 | chr18 | + | 12409891 | 12409946 | GCACTCCGGCCTGGGTGACAGAGCAAGACTCCGTCTCAAACAAAAAAGTCATTCAG |
| *AFAP1L1* | 84006 | 4974 | 5A | 50 | chr5 | + | 148664027 | 148664076 | CTGATGGACCTTGGGCTGATTGCACCTTTTGGCTATGGTGCTTGGATCAG |
| *Not prioritised* | | | | | | | | | |
| *NUCB1* | 274872 | 16698 | 8A' | 33 | chr19 | + | 54114147 | 54114179 | ATGGAGGAGGAGCGACTGCGCATGCGGGAGCAT |
| *TIA1* | 51127 | 3043 | 4A | 33 | chr2 | - | 70309695 | 70309727 | GTAGTACCGTTGTCAGCACACAGCGTTCACAAG |
| *TNXB* | 116531 | 7005 | U3 | 38 | chr6 | - | 32203238 | 32203275 | TTTGACGGCAGCTCCCTGGACGTGGGGATGGATGTCAG |
| *SFI1* | 299986 | 18375 | U1 | 42 | chr22 | + | 30152727 | 30152768 | CCTATTGGAAAAGATCTGGGACTATCTGAAACTAGTGAGAAT |
| *OVOL2* | 287577 | 17608 | W2 | 59 | chr20 | - | 17885623 | 17885681 | CTGAGATCTGTATCTGTGGACCTGAATGTTGATCCCTCGCTTCAGATTGACATACCTGA |
| *RAD51L1* | 214776 | 13080 | 10A | 45 | chr14 | + | 68005025 | 68005069 | ATCCTATGGCATGAAGTGGGCAATGATTCTGACACCAGTGTGGAG |
| *MX1* | 293019 | 17978 | U4 | 45 | chr21 | + | 41715126 | 41715170 | GACGGGCAGGAGACAGATGCCTTCCTCTTGTCTCAACTGCAAGAG |
| *MPV17L* | 235978 | 14243 | W1 | 46 | chr16 | + | 15516663 | 15516708 | ATCAGCTGGACATCATCTCCATGGCGGAGACAACCATGATGCCAGA |
| *WASF1* | 118141 | 7091 | 2A | 57 | chr6 | - | 110605643 | 110605699 | CTTGTTGCATTTACCCTTTTGATAAAAGAGAATCACATAAGCATTGCAGAGGCAGTT |
| *ALDH3B2* | 168876 | 10159 | U1 | 49 | chr11 | - | 67205223 | 67205271 | CTGCCACCATGTGAAGAAGGATGTGGTTGCTTCCCCTTCCACCATAACT |
| *SIRT3* | 182299 | 11146 | 2b' | 49 | chr11 | - | 223438 | 223486 | TGTTGTTGGAAGTGGAGGCAGCAGTGACAAGGGGAAGCTTTCCCTGCAG |
| *MTRF1* | 205777 | 12538 | 1B | 71 | chr13 | - | 40734547 | 40734617 | GAAAGTATTCCATGCTACATGTGCAAGAAACTTGGAAAACACTGAAAAGTAGAAAAA ATAGCAAGCAAAAG |
| *CHID1* | 171768 | 10341 | 2A | 50 | chr11 | - | 893704 | 893753 | GCTGGAGCGCAATGGCGCGATCTTGGCTCACCGCAACCTCTGCCACCCAG |
| *TJAP1* | 116322 | 6990 | 2a' | 50 | chr6 | + | 43553803 | 43553852 | GCTTAGATCAGCCTTTCCACAGCTGTTAGCAGCATCTGCCCCAATTTCAG |
| *DNAJC4* | 173081 | 10415 | 4c' | 80 | chr11 | + | 63756478 | 63756557 | CTGCACCCAGACCGGGACCCTGGGAACCCAAGCCTGCACAGCCGCTTTGTGGAGCTGA GCGAGGCATACCGTGTGCTCAG |
| *BMP4* | 208853 | 12715 | 3a' | 53 | chr14 | - | 53488645 | 53488697 | AGACACCATGATTCCTGGTAACCGAATGCTGATGGTCGTTTTATTATGCCAAG |
| *TIAM2* | 116268 | 6988 | U6 | 95 | chr6 | + | 155376748 | 155376842 | ACACACACACACACACACACACACACACACACCTGAGATGGGGTAGATCATTGTATTT TTGTGTCTACCAGCAAGAAAAGGAAGGAAAAACTAAG |
| *MTP18* | 298212 | 18272 | 3b' | 69 | chr22 | + | 29153158 | 29153226 | GTGCCCAGCCCTGAAGCAGGCCGCAGCGCCAGGGTGACTGTGGCTGTGGTGGACACC TTTGTATGGCAG |
| *SEZ6* | 255662 | 15470 | 8a' | 55 | chr17 | - | 24310982 | 24311036 | CCTTCCAGCAGGGCCATTGCTATGAGCCCTTTGTCAAATACGGTAACTTCAGCAG |

**The 24 exons identified only in cycloheximide-treated HMEC, with >20 reads.** Chr, chromosome. Start and end coordinates as originally aligned to GRCh18.
